# Inhibition of SAR S-CoV-2 infection and replication by lactoferrin, MUC1 and α-lactalbumin identified in human breastmilk

**DOI:** 10.1101/2021.10.29.466402

**Authors:** Xinyuan Lai, Yanying Yu, Wei Xian, Fei Ye, Xiaohui Ju, Yuqian Luo, Huijun Dong, Yihua Zhou, Wenjie Tan, Hui Zhuang, Tong Li, Xiaoyun Liu, Qiang Ding, Kuanhui Xiang

## Abstract

The global pandemic of COVID-19 caused by the severe acute respiratory syndrome coronavirus-2 (SARS-CoV-2) infection confers great threat to the public health. Human breastmilk is an extremely complex with nutritional composition to nourish infants and protect them from different kinds of infection diseases and also SARS-CoV-2 infection. Previous studies have found that breastmilk exhibited potent antiviral activity against SARS-CoV-2 infection. However, it is still unknown which component(s) in the breastmilk is responsible for its antiviral activity. Here, we identified Lactoferrin (LF), MUC1 and α-Lactalbumin (α-LA) from human breastmilk by liquid chromatography-tandem mass spectrometry (LC-MS/MS) and in vitro confirmation that inhibited SARS-CoV-2 infection and analyzed their antiviral activity using the SARS-CoV-2 pseudovirus system and transcription and replication-competent SARS-CoV-2 virus-like-particles (trVLP) in the Huh7.5, Vero E6 and Caco-2-N cell lines. Additionally, we found that LF and MUC1 could inhibit viral attachment, entry and post-entry replication, while α-LA just inhibit viral attachment and entry. Importantly, LF, MUC1 and α-LA possess potent antiviral activities towards not only wild-type but also variants such as B.1.1.7 (alpha), B.1.351 (beta), P.1 (gamma) and B.1.617.1 (kappa). Moreover, LF from other species (e.g., bovine and goat) is still capable of blocking viral attachment to cellular heparan sulfate. Taken together, our study provided the first line of evidence that human breastmilk components (LF, MUC1 and α-LA) are promising therapeutic candidates warranting further development or treatingVID-19 given their exceedingly safety levels.

## Introduction

It confers a great threat to global public health since the pandemic of coronavirus disease 2019 (COVID-19) caused by severe acute respiratory syndrome coronavirus 2 (SARS-CoV-2) since outbreak in the late of 2019, which showed close phylogenetic relationship with SARS-CoV outbreak in 2002 ^1^. The high rates of human deaths and pandemic distribution worldwide have caused the collapse of health systems in many countries, especially in developing countries^2^. In addition, the ongoing pandemic has resulted in several viral variants^3-5^. These variants may potentially alter viral transmission, pathogenicity, efficacy of drug treatment as well as the binding capacity of vaccine elicited antibodies^6-9^. In this situation, it raises the need of several effective drugs with low toxicity to treat this disease.

As reported previously, COVID-19 pandemic also confers great concern of mother-to-child transmission (MTCT) by breastfeeding^10-14^. In addition to the report of SARS-CoV-2 RNA was detected in human breastmilk, it was still unclear that SARS-CoV-2 could transmit to infants through breastfeeding^13^. Several evidences in the clinic study showed that SARS-CoV-2 couldn’t transmit to infants by breastfeeding^10-12,15,16^. However, it is controversial if live SARS-CoV-2 existing in the breastmilk could still be infectious.

Human milk is uniquely suited to breastfeed human infants due to its nutritional composition and bioactive factors for promoting antimicrobial and immunomodulatory effects^17^. Breastmilk could inhibit several viruses infection, such as human immunodeficiency virus (HIV), cytomegalovirus (CMV) and dengue virus. It was reported that breastmilk could not only inhibit several enveloped viruses such as herpes simplex types 1 and 2, HIV and CMV, but also showed effective activity in vitro against many non-enveloped viruses like rotavirus, enterovirus and adenovirus^17,18^. The lactoferrin (LF), a important component in breastmilk, could suppress SARS-CoV and SARS-CoV-2 through blocking virus to bind heparan sulfate proteoglycans, raising a concern that human breastmilk can also suppress SARS-CoV-2 infection^19-21^. Our previous study had confirmed that human breastmilk significantly inhibits SARS-CoV-2 and its related pangolin coronavirus (GX_P2V) infection and have strong potent anti-SARS-CoV-2 effects than LF treatment alone, indicating that there are potential other factors in breastmilk also play important role in inhibiting SARS-CoV-2 inhibition and replication^22^. Thus, it is still unclear that which components are potential to suppress SARS-CoV-2 infection in an overall study.

Breastmilk is an extremely complex to nourish infants and protect them from different kinds of disease. It contains more than 400 different proteins and many of them exhibit the antimicrobial activity^23^. Proteins in human milk can be divided into three groups, caseins, mucin and whey proteins. These bioactive proteins from the whey fraction include LF, mucins (MUC1and MUC4), α-lactalbumin (α-LA), lactadherin, lactoperoxidase, IgA and lysozyme (LZ) and so on^23,24^. LF is an iron-binding protein with two molecules of Fe^3+^ per protein and rich in human milk (1-2 g/L in mature milk, 5-10 g/L in colostrum), which relatively lower in other species such as bovine milk (0.02-0.3g/L in mature milk, 2-5 g/L in colostrum)^24^. As reported previously, LF is an important component to protect infants from microbes and virus infection^17,24^. LZ is also known to be a key protein inhibiting bacteria and rich in human milk (0.4g/L). Lactadherin, a 46 KDa mucin associated protein, was reported that it displayed high viral receptor binding and showed specific anti-rotavirus activity^24^. MUC1 has the antiviral activity to inhibit influenza virus infection^25^. In addition, human milk also contains antibodies like IgG and IgA which show antiviral activity^24,26^. Therefore, these gave us confidence to continue to identify the anti-SARS-CoV-2 components in breastmilk and explore the underlying mechanisms.

Here, we aimed to identify the protein components in human breastmilk by the liquid chromatography-tandem mass spectrometry (LC-MS/MS) ^27^ and verify their antiviral activity using the SARS-CoV-2 pseudovirus and trVLP system in Huh7.5, Vero E6 and Caco-2-N cell lines, respectively^28^. We identified that LF, MUC1 and α-LA could inhibit SARS-CoV-2 and variants infection by blocking viral entry and post-entry replication, which shed light on the important role of LF, MUC1 and α-LA derived from human milk and provide the clues for the development of antiviral drug identification and design.

## Results

### LF, MUC1 and α-LA were identified in skimmed milk with inhibition of SARS-CoV-2 infection

To identify the effective factors of human skimmed milk influencing on SARS-CoV-2 infection, we performed mass spectrometry to identify the potential factors from the skimmed milk shown in **Figure 1A**. The skimmed milk was separated by cation exchange column, anion exchange column and size exclusion column in sequence. After these column separations, we got several fractions such as SP1, SP2, SP3, Q1 and Q2, which were subsequently collected for anti-SARS-CoV-2 analysis (**Figure 1B to1D**). We infected the Vero E6 cells with 650 TCID_50_/well of SASRS-CoV-2 pseudovirus and added same ratio of SP1, SP2, SP3, Q1 and Q2 into the media, the skimmed breastmilk (A17) was used as control. One day post infection, we found that SP2, SP3 and Q2 exhibited inhibitory effects to SARS-CoV-2 pseudovirus infection (**Figure 1E**). Then, we selected these fractions into tandem mass spectrometry (MS/MS) analysis and found that LF were the potential factors for SARS-CoV-2 inhibition (**Figure 1F and Figure S1 to S4**). These data suggested that LF is the potential factor in skimmed milk inhibiting SARS-CoV-2 infection.

**Fig. 1.**
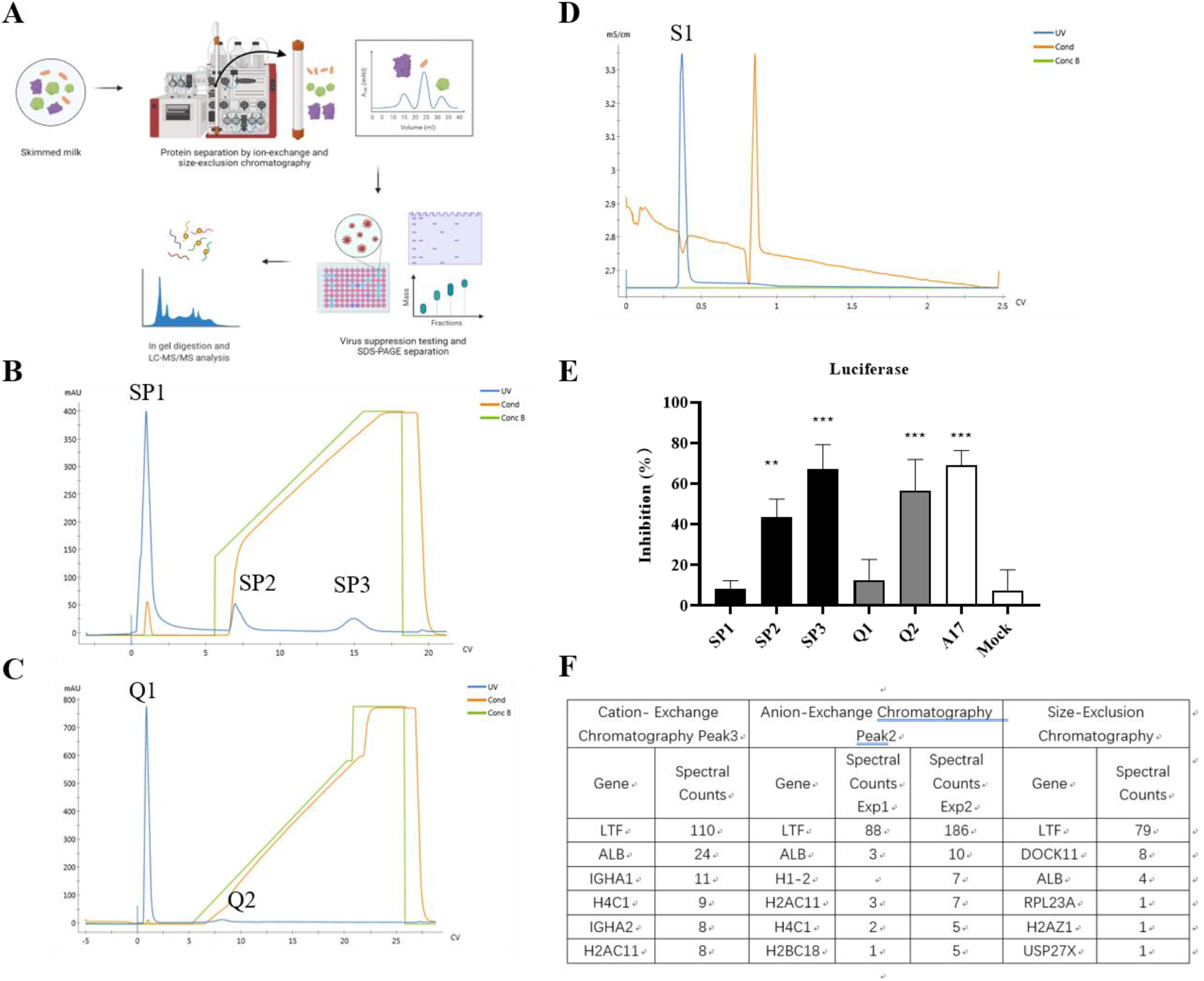
Identification of the key factors from skimmed milk for inhibiting SARS-CoV-2 infection by mass spectrometry. (A) Schematic illustration of identification of key factors from skimmed milk by mass spectrometry. The skimmed milk was separated through the cation exchange column (B), the anion exchange column (C), and the size exclusion column (D) in sequence. The UV signal indicates the protein fraction (280 nm). (E) Inhibition analysis of the collected fractions to the inhibition of SARS-CoV-2 pseudovirus infection. The Vero E6 cells were infected with 650 TCID50/well of SARS-CoV-2 pseudovirus and treated with different fractions with same ration at the same time. The human skimmed breastmilk (A17) was used as positive control. The cells were lysed and the luminescence was measured according to the manufacture’s instruction 24 hours post-infection. (F) Table with top 6 (based on number of spectral counts) hits from skimmed milk in eluate fractions determined by tandem mass spectrometry (MS/MS). Data are presented as mean± SD and repeated at least three times (N = 3), \*\**p* < 0.01, \*\*\**p* < 0.001.

As reported in our previous publication, LF might not be the effective factor alone in breastmilk and there might be other components involving the anti-SARS-CoV-2 infection^22^. Previous reports showed that some factors, such as MUC1, MUC4, lactadherin, α-LA, etc., have anti-microbe activity^23,24^. To identify which of them could also inhibit SARS-CoV-2 infection, we mixed the MUC4, MUC1, α-LA, lactadherin and LF with 650 TCID_50_/well of SASRS-CoV-2 pseudovirus with luciferase expression, respectively. Then, the mixture was added to infect Vero E6 cell for 24h. MUC4 (**Figure 2A**) and lactadherin (**Figure 2D**) were showed no suppression to SARS-CoV-2 pseudovirus infection. Interesting, MUC1 (0.25 μg/ml, **Figure 2B**), α-LA (0.25 mg/ml, **Figure 2C**) and LF (0.25 mg/ml, **Figure 2E**) showed dramatic inhibition of SARS-CoV-2 pseudovirus infection. In addition, we also used the GFP expressing SARS-CoV-2 pseudovirus to verify the inhibition activity of LF, MUC1 and α-LA in both Huh7.5 and Vero E6 cell lines. As shown in **Figure 2F to 2H**, all the LF, MUC1 and α-LA could suppress GFP expression, indicating that these factors in skimmed milk could inhibit SARS-CoV-2 infection. Thus, these three factors in breastmilk were validated to inhibit SARS-CoV-2 pseudovirus infection.

**Fig. 2.**
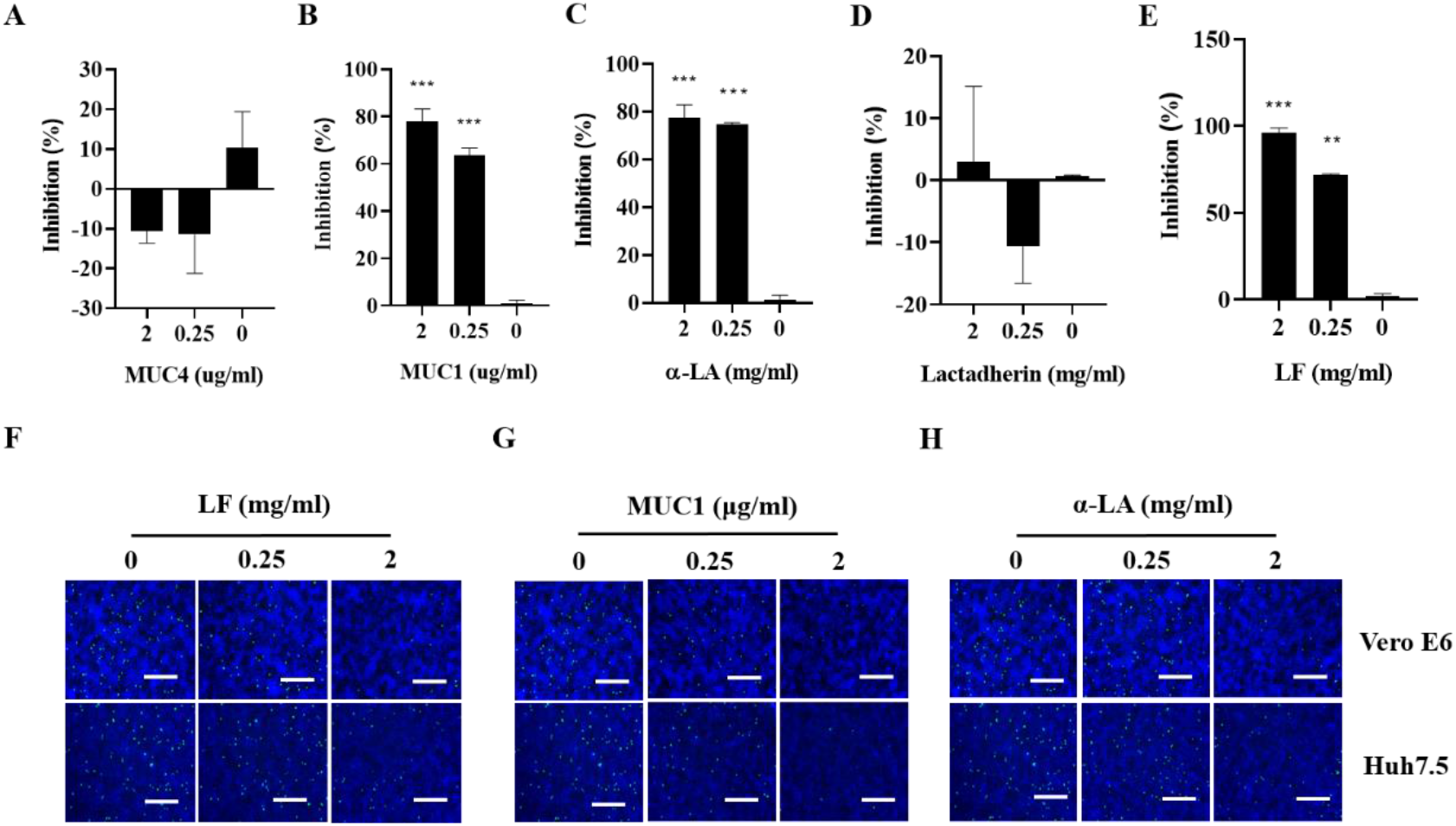
Identification of the key factors from the reported skimmed milk’s factors for inhibiting SARS-CoV-2 infection. Inhibition analysis of MUC4 (A), (MUC1), α-LA (C), lactodherin (D) and LF (E) to SARS-CoV-2 pseudovirus infection. The Vero E6 cells were infected with 650 TCID_50_/cell of SARS-CoV-2 pseudovirus and treated with different concentration of MUC4, MUC1, α-LA, lactadherin and LF at the same time. The cells were lysed and the luminescence was detected according to the manufacture’s instruction 24 hours post-infection. Fluorescence analysis of different concentration of LF (F), MUC1 (G) and α-LA (H) to inhibit SASR-CoV-2 pesudovirus with GFP expression in both Vero E6 and Huh 7.5 cells. Scale bar represents 100 μm. α-LA, a-lactalbumin; LF, lactoferrin. Data are presented as mean± SD and repeated at least three times (N = 3), \*\**p* < 0.01, \*\*\**p* < 0.001.

### LF, MUC1 and α-LA suppress SARS-CoV-2 infection and replication in the trVLP infected Caco-2-N cell system

To further explore these factors inhibition ability of SARS-CoV-2 infection, we evaluated the effects of LF, MUC1 and α-LA on SARS-CoV-2 infection in the cell culture system of transcription and replication-competent SARS-CoV-2 virus-like-particles (trVLP), which expresses a reporter gene (GFP) replacing viral nucleocapsid gene (N) and also could complete viral life cycle in the cells expression N proteins^28^. Caco-2-N cells were infected with trVLP (MOI=1) and treated with LF, MUC1 and α-LA at different concentration for 96h (**Figure 3A**). After the treatment, RT-qPCR analysis of cell lysates showed a dramatic decrease in the levels of SARS-CoV-2 RNA (**Figure 3B**). Similarly, GFP expressing reflecting the viral replication revealed that both LF and MUC1 could significantly suppress SARS-CoV-2 replication with low GFP expression (**Figure 3C**). Although we could see the inhibition effect of α-LA on SARS-CoV-2 infection, the inhibitory ability is relatively lower, which still showed a few GFP expression even in concentration of 2 mg/ml. Consistent to the GFP results, western blot analysis of the cell lysates showed that MUC1 and LF could significantly suppress the S protein expression of SARS-CoV-2 (**Figure 3D**). The α-LA at high concentration of 2 mg/ml could also suppress SARS-CoV-2 infection.

**Fig. 3.**
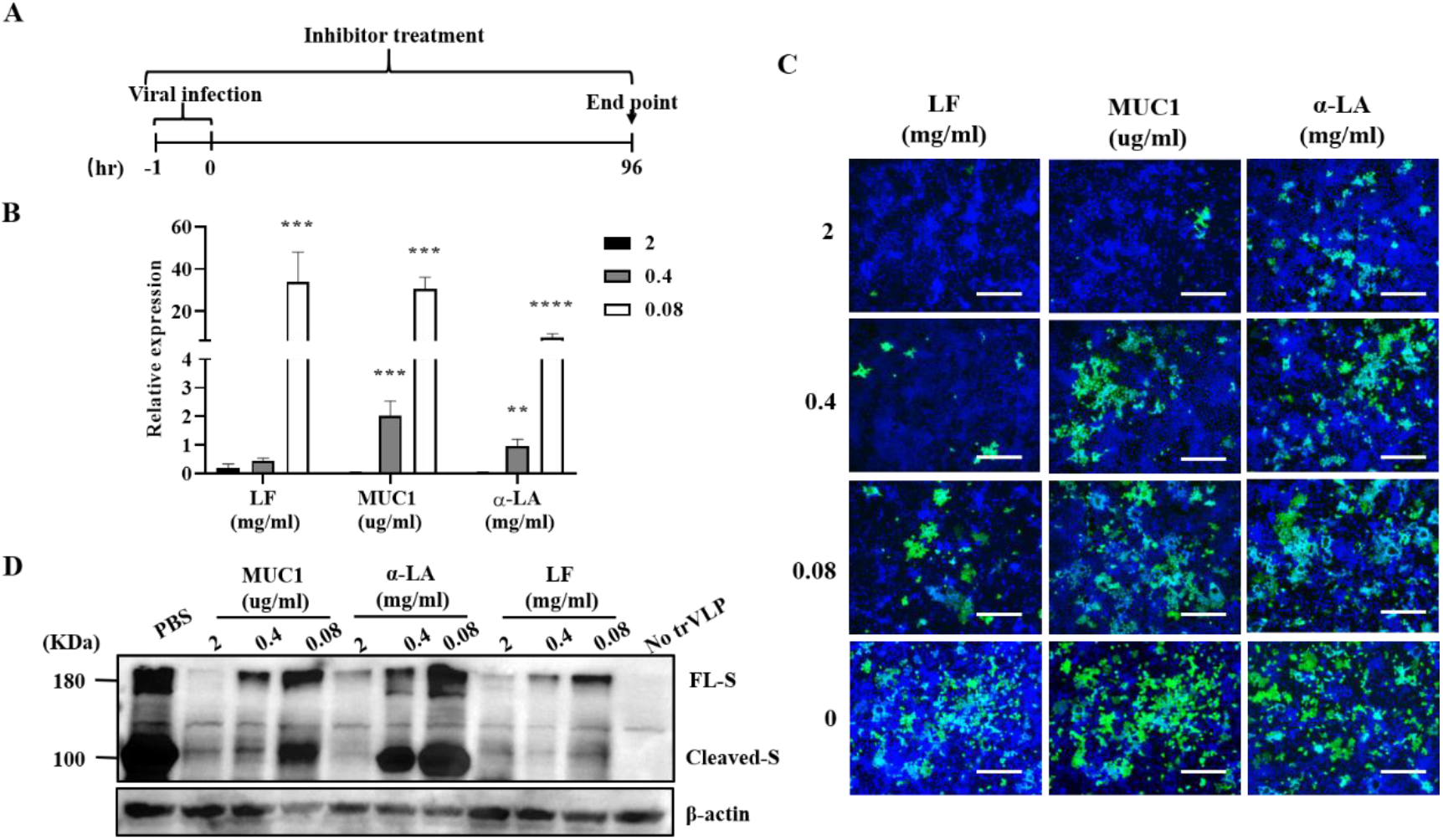
MUC1, a-lactalbumin and lactoferrin suppress viral infection and replication in SARS-CoV-2 infected Caco-2-N cells. (A) Schematic illustration of the experiment. (B) RT-qPCR analysis, (C) fluorescence analysis and (D) western blot analysis were used to detect viral RNA, GFP and viral spike protein in SARS-CoV-2 infected Caco-2-N cells treated with different doses of MUC1, α-LA and LTF. Caco-2-N cells were infected with trVLP (SARS-CoV-2 N protein deficient particles, MOI=1) and treated with different doses of LF, MUC1 and α-LA. α-LA, a-lactalbumin; LF, lactoferrin. Scale bar represents 100 μm. Data are presented as mean± SD and repeated at least three times (N = 3), \*\**p* < 0.01, \*\*\**p* < 0.001, \*\*\*\**p*<0.0001.

In addition, to explore whether the LF from different species possessed inhibitory effect on SARS-CoV-2 infection, we tested the inhibitory effect of recombinant human LF (rLF), human isolated LF (hLF), bovine LF (bLF) and goat LF (gLF) on SARS-CoV-2 infection. The skimmed milk (A17) was used as positive control. Firstly, we tested them in the SARS-CoV-2 pseudovirus system. We infected the Vero E6 cells with 650 TCID_50_/well of SASRS-CoV-2 pseudovirus and treated the cells with rLF, hLF, bLF, gLF and A17 for 24h. After treatment, luciferase assay showed that all of the LFs and A17 could significantly suppress SARS-CoV-2 pseudovirus infection, indicating that LF from bovine and goat could also inhibit SASR-CoV-2 infection (**Figure 4A**). Consistent to the pseudovirus system, trVLP system showed that all the LFs could also inhibit SARS-CoV-2 infection and replication in RT-qPCR analysis (**Figure 4B**) and immune microscopy analysis of GFP expression (**Figure 4C**). Western blot analysis of the cell lysates showed that rLF and bLF significantly suppressed SARS-CoV-2 S protein expression (**Figure 4D**). In addition, we found that gLF at high concentration of 2 mg/ml could be toxic to cell (**Figure 4E**), while other LFs showed no toxic to cell. When treating the cells at different dosages and infecting the cells with trVLP (MOI=1) for 96 h, we found that gLF suppressed SARS-CoV-2 infection in a dose dependent manner. However, CCK8 analysis showed that gLF at high concentrations exhibited high cytotoxicity to Caco-2-N cells with CC_50_ of 5.64 mg/ml (**Figure 4F**). Thus, LF, MUC1 and α-LA from human breastmilk were confirmed to suppress SARS-CoV-2 infection and replication. In addition, LF from other species could also inhibit SARS-CoV-2 infection.

**Fig. 4.**
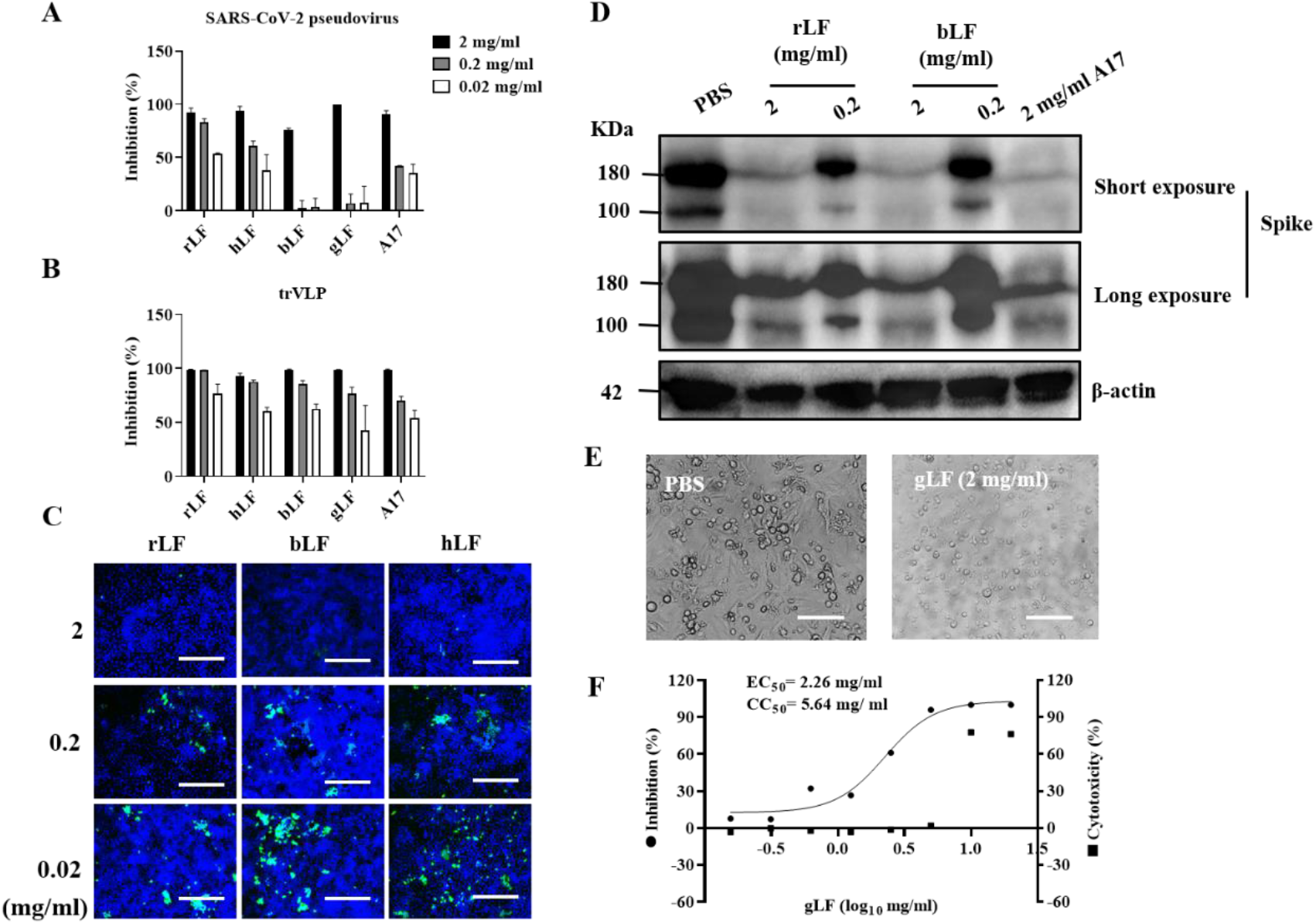
LF from different species also suppressed SARS-CoV-2 infection and replication. (A) Inhibition analysis of recombinant LF (rLTF), human LF (hLF), bovine LF (bLF) and goat LF (gLF) to SARS-CoV-2 pseudovirus infection in Vero-E6 cells. Skimmed milk (A17) was used as positive control for SARS-CoV-2 inhibition. RT-qPCR (B), immunoplot (C) and western blot (D) were used to detected viral RNA, GFP and viral spike protein in trVLP infected Caco-2-N cells treated with different doses of rLF, hLF, bLF, gLF and A17. (E) High concentration of gLF are toxic to cells in the Vero E6 cell model. (F) The gLF treatment dose-dependently suppressed viral infection and replication, but with cell toxicity at high doses, in trVLP infected Caco-2-N cells. Scale bar represents 100 μm. Data are presented as mean± SD and repeated at least three times (N = 3), \*\**p* < 0.01, \*\*\**p* < 0.001, \*\*\*\**p*<0.0001.

### LF, MUC1 and α-LA dose and time dependently decrease SARS-CoV-2 infection and replication in association with reduced viral RNA

We performed a dose-response experiment with LF, MUC1 and α-LA to assess their suppression on SARS-CoV-2 infection and replication. Firstly, we explored them in the trVLP system infected Caco-2-N cells. The results confirmed that the inhibitory effects of MUC1, LF and α-LA on SARS-CoV-2 infection were dose dependent and the 50% effective concentration (EC_50_) for MUC1, LF and α-LA was as low as 0.04 μg/ml (**Figure 5A**), 0.08 mg/ml (**Figure 5B**) and 0.68 mg/ml (**Figure 5C**), respectively. Surprisingly, the cytotoxic values evaluated by CCK-8 assay showed that MUC1, LF and α-LA have no cytotoxicity to the cells, even the highest concentration of MUC1(2 μg/ml), LF (2 mg/ml) and α-LA (2 mg/ml). Consistent to these results, we also tested them in the SARS-CoV-2 pseudovirus system infected Vero E6 cells. As shown in **Figure S5A**, the results confirmed the inhibitory effects of MUC1 on SARS-CoV-2 pseudovirus infection and exhibited that the EC_50_ for MUC1 was as low as 0.1 μg/ml. Similarly, the EC_50_ of LF and α-LA was as low as 0.1 mg/ml (**Figure S5B**) and 0.16mg/ml (**Figure S5C**), respectively.

**Fig. 5.**
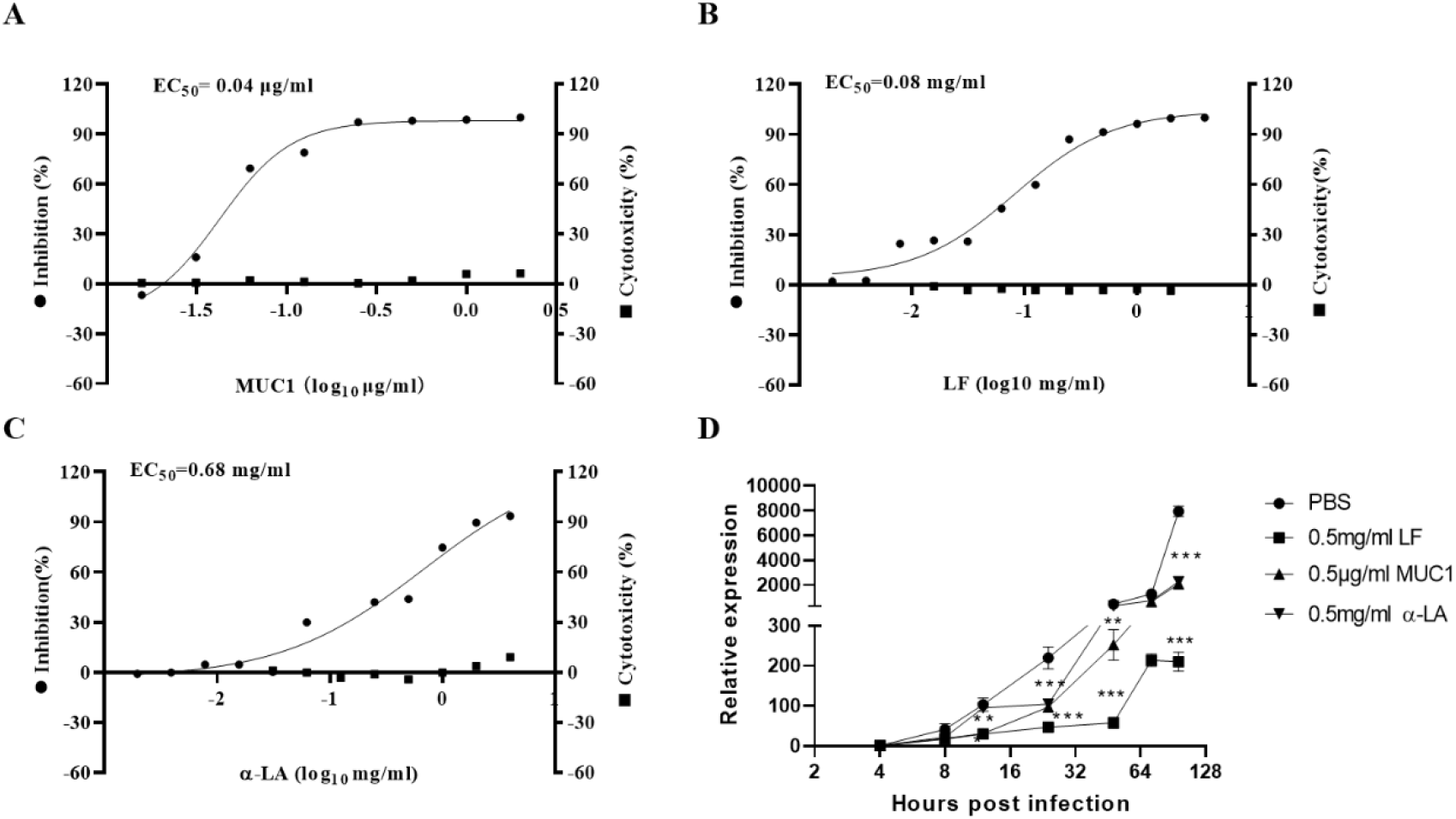
MUC1, LF and α-LA treatment dose-dependently suppressed viral infection and replication with reduced viral RNA in trVLP infected Caoco-2-N cells. The inhibition analysis reflected by intracellular viral RNA of trVLP infected Caco-2-N cells treated with serial doses of MUC1 (A), LF (B)and α-LA (C) were determined by RT-qPCR. (D) The viral RNA levels with time dependent suppressed by MUC1, LF and α-LA were detected by RT-qPCR. The Caco-2-N cells were infected with trVLP (MOI=1) and treated with MUC1, LF and α-LA at the same time, then the cells were harvested at different timepoints as designed. The cells were lysed and detected by RT-qPCR. Data are presented as mean± SD and repeated at least three times (N = 3), \*\*\**p* < 0.001.

When treating the cells with relative lower concentration of MUC1, LF and α-LA in the trVLP infected Caco-2N cells, we found that LF and MUC1 could significantly suppressed SARS-CoV-2 infection at early time point and keep the inhibitory effects during the culture (**Figure 5D**). However, although α-LA inhibited SARS-CoV-2 infection at early time point, the inhibitory ability decreased during the culture, suggesting that α-LA might be the inhibitor influence on the early steps of viral life cycle, such as attachment and entry.

### LF and MUC1 affect viral attachment, entry and post-entry of viral replication, but α-LA decreases the viral attachment and entry

To explore which stage of SARS-CoV-2 replication cycle targeted by LF, MUC1 and α-LA, we treated the Caco-2-N cells with individual factors at different time point during viral infection. The cells were treated with these factors before and after viral infection. Firstly, we designed the attachment experiments as shown in **Figure 6A** (pre-infection treatment). In the pre-infection treatment experiments, these factors were washed away before viral infection. The levels of viral RNA were significantly lower in those group treated by MUC1, α-LA, rLF, hLF, bLF and A17 than those treated by PBS (**Figure 6B**). Similarly, the GFP expressing results showed that cells treated with these factors nearly expressed no GFP (**Figure 6C**). We also tested them in the pseudovirus system (**Figure S6A**). Consistent reductions in the luciferase expression were identified in the SARS-CoV-2 pseudovirus infected Vero E6 and Huh7.5 cells (**Figure S6B**). These data indicated that MUC1, LF and α-LA might attach the cells to block viral attachment and entry.

**Fig. 6.**
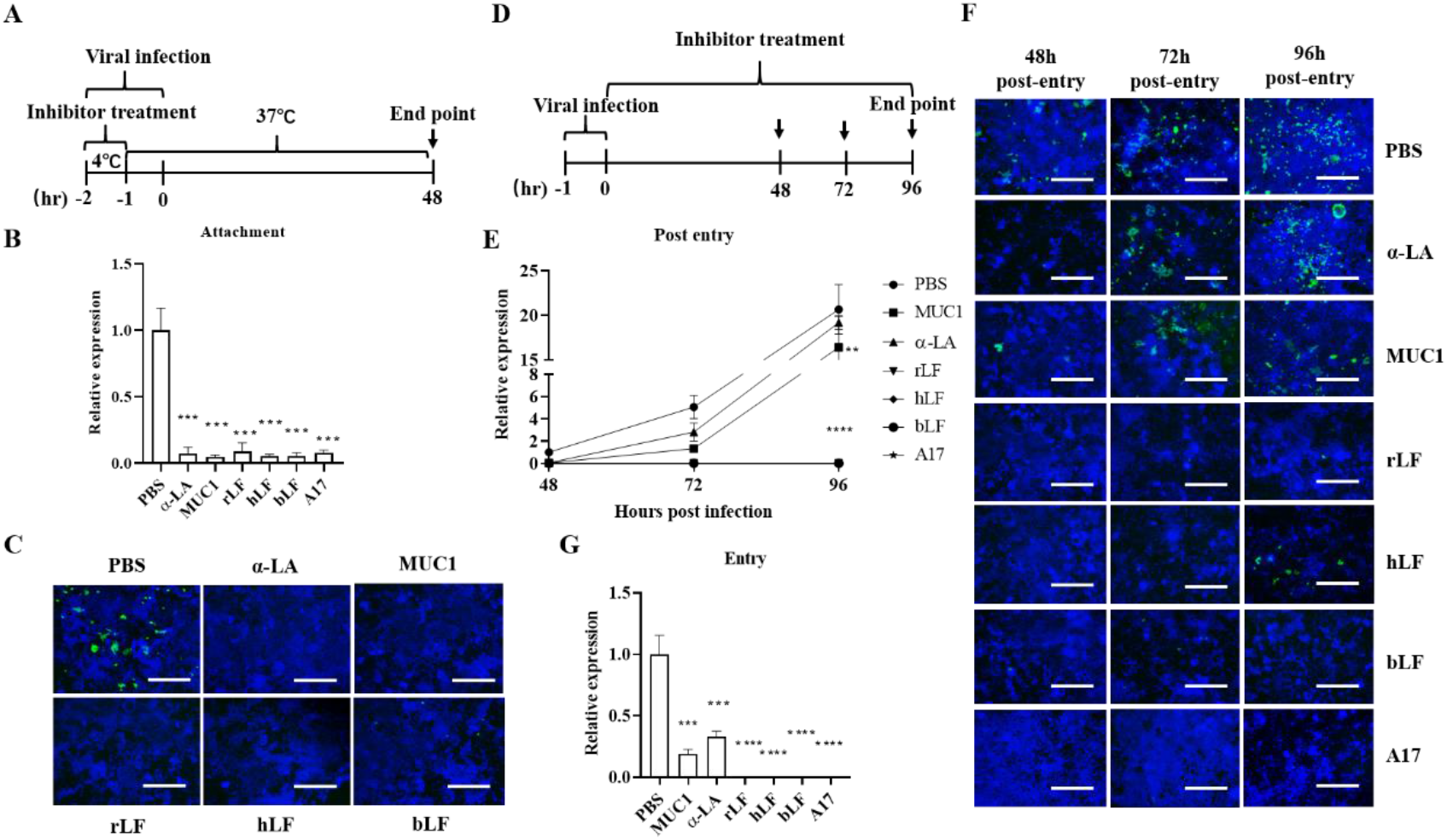
MUC1, LF and α-LA treatment affects the early stage of viral infection including attachment and entry, but LF also decreases viral RNA levels after virus entry. (A) Schematic illustration of the pretreatment experiment. RT-qPCR (B) and immune blot (C) assays were used to detected viral RNA and GFP expression in trVLP infected Caco-2-N cells. Skimmed milk (A17) was used as positive control for inhibiting SARS-CoV-2 infection. (D) schematic illustration of the post-infection treatment experiments. RT-qPCR (E) and immune blot (F) assays were used to detected viral RNA and GFP expression at different time points in trVLP infected Caco-2-N cells. (G) The viral RNA levels detected by RT-qPCR are suppressed by MUC1, α-LA, rLF, hLF, bLF and A17 during viral entry. The trVLP was mixed with Caco-2-N cells at 4 °C for 1 hour then discarded the supernatant and washed with PBS for three times. The cells were then added with fresh media with MUC1, α-LA, rLF, hLF, bLF and A17 treatment at 37 °C for 2 hours. After that, the supernatant was disregarded and the cells were reloaded with fresh media and cultured for 48 hours. The cells were lysed and the viral RNA was detected by RT-qPCR. MUC1 (2 ug/ml), α-LA (a-lactalbumin, 2 mg/ml), rLF (recombinant lactoferrin, 2 mg/ml), hLF (human lactoferrin, 2 mg/ml), bLF (bovine lactoferrin, 2 mg/ml) and A17 (skimmed milk, 2 mg/ml). Scale bar represents 100 μm. Data are presented as mean± SD and repeated at least three times (N = 3), \*\*\**p* < 0.001, \*\*\*\**p*<0.0001.

Then, we also designed the post-infection experiment as shown in **Figure 6D**. In the post-infection treatment experiments, the viral RNA and GFP expression were reduced by treatment with MUC1 and LF, while α-LA treatment resulted in nearly no reduction of RNA and GFP expression (**Figure 6E** and **6F**). Consistent reductions in the luciferase expression of pseudovirus system (**Figure S6C**) after MUC1 and LF treatment and less reduction by α-LA treatment in the SARS-CoV-2 pseudovirus (**Figure S6D**). The pseudovirus was packed based on the VSV. Thus, the post-entry replication usually reflects VSV replication. Then, we applied the VSV as control to performed the experiment as designed in **Figure S6C**. Expectedly, MUC1, α-LA and LF also inhibited VSV replication (**Figure S6E**). These data suggested that LF and MUC1 suppress SARS-CoV-2 post-entry replication. α-LA has limited suppression on SARS-CoV-2 post-entry replication.

In addition, we also performed the experiment to explore if MUC1, LF and α-LA suppress viral entry. We infected trVLP into the Caco-2-N cells and pseudovirus into the Huh7.5 and Vero E6 cells at 4°C for 1h, respectively. Then, we washed the virus away and changed the media supplemented with these factors until the end point of infection. Interestingly, all these factors showed potent antiviral activity as evidence by the dramatic reduction of the SARS-CoV-2 RNA (**Figure 6G**) and luciferase activity levels (**Figure S6F**). These data suggested that MUC1, LF and α-LA suppress SARS-CoV-2 entry step of viral life cycle.

### The α-LA interferes the interaction of ACE-2 and S protein, MUC1 and LF suppresses SARS-CoV-2 attaching to the HSPG

To explore if MUC1, LF and α-LA interfere the interaction of ACE-2 and S protein, we performed the affinity assay. As shown in **Figure 7A** and **7B**, LF and MUC1 didn’t block the interaction of ACE-2 and S protein, indicating that LF and MUC1 suppress SARS-CoV-2 infection through other ways rather than the ACE-2target. Surprisingly, α-LA could interfere SARS-CoV-2 attachment to ACE-2, showing high absorbance after α-LA treatment at low concentration of 0.25 mg/ml (**Figure 7C**).

**Fig. 7.**
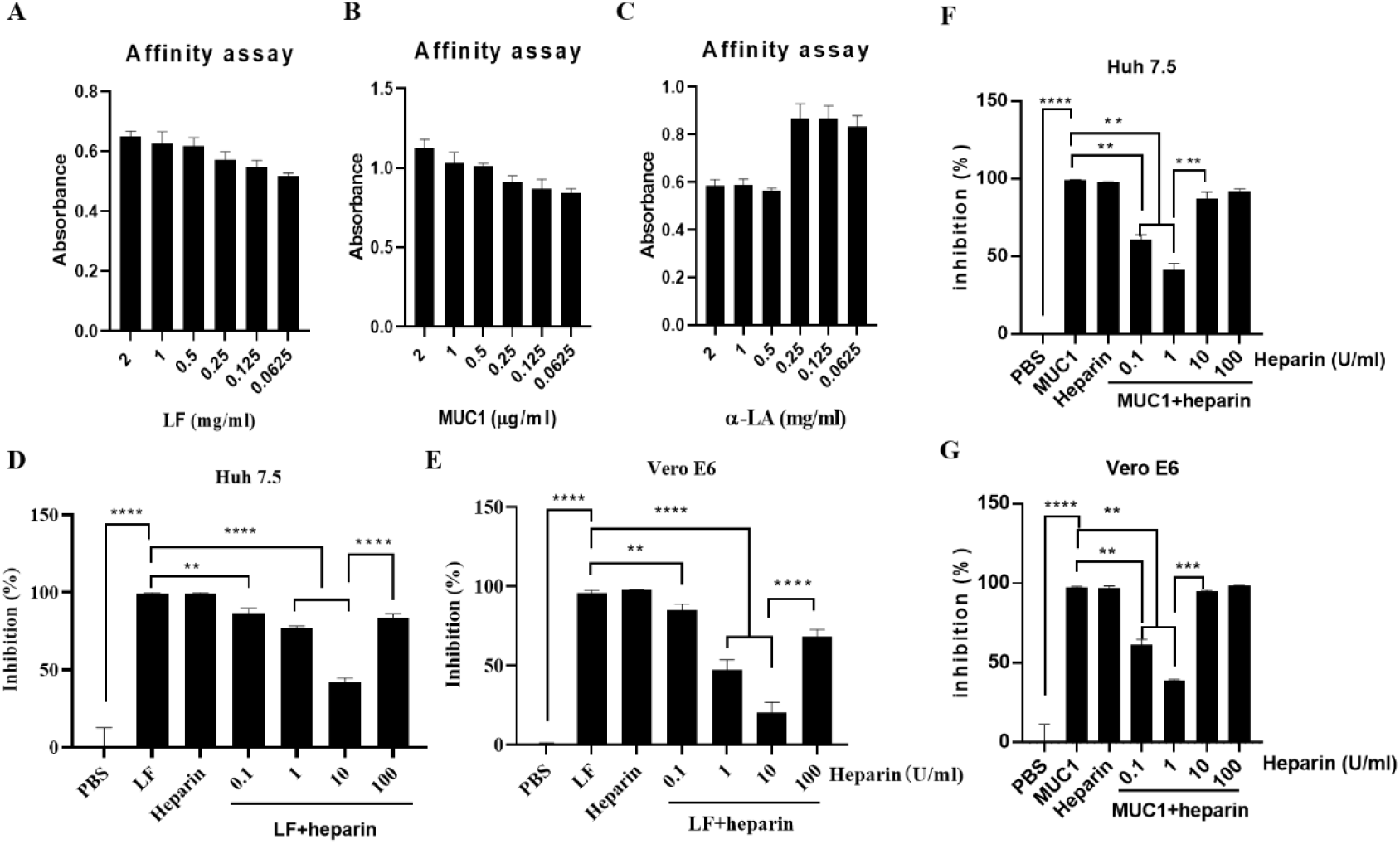
α-LA interferes the affinity of spike protein and ACE-2, MUC1 and LF inhibits SARS-CoV-2 attaching to HSPG. Affinity assay for the impact of LF (A), MUC1 (B) and α-LA (C) on the affinity between ACE2 and SARS-CoV-2 RBD. (D) and (E) inhibition of SARS-CoV-2 pseudovirus infection of Huh7.5 and Vero E6 cells were treated with heparin for 10 min at the dosage of 0.1 U/ml, 1 U/ml, 10 U/ml and 100 U/ml. Then, 2 mg/ml LF was added to each group and incubated at 37°C for 1h. Similarly, (F) and (G) inhibition of SARS-CoV-2 pseudovirus infection of Huh7.5 and Vero E6 cells were treated with heparin for 10 min at the dosage of 0.1 U/ml, 1 U/ml, 10 U/ml and 100 U/ml. Then, 2 mg/ml LF was added to each group and incubated at 37°C for 1h. The luciferase assay was performed to detect SARS-CoV-2 pseudovirus infection. Data are presented as mean± SD and repeated at least three times (N = 3), \*\**p* < 0.01, \*\*\*\**p*<0.0001. HSPG: heparan sulfate proteoglycans; LF: lactoferrin; MUC1: mucin 1.

Moreover, it was reported that LF inhibit SARS-CoV infection through the HSPG target^19^. We also performed the experiment to explore the possible same targetfor LF to inhibit SARS-CoV-2 infection. We treated the cells with LF combined with different concentrations of heparin. We found that the inhibitory effect of LF and MUC1 on SARS-CoV-2 decreased with dose-dependent of heparin combined treatment, to the deep when 10 U/ml of heparin was added in both Huh7.5 (**Figure 7D and 7F**) and Vero E6 (**Figure 7E and 7G**) cells. However, when increasing the heparin to 100 U/ml, we found that the inhibitory effect increased, indicating that heparin could interfere the inhibitory effect of LF on SARS-CoV-2 infection.

To further confirm that LF and MUC1 interact SARS-CoV-2 attaching HSPG, we also performed new experiments in the trVLP infected Caco-2-N cells as designed in **Figure S7A**. as the results, we also found that MUC1 (**Figure S7B**) and LF (**Figure S7C**) rather than α-LA (**Figure S7D**) could interfere heparin’s inhibition of SARS-CoV-2 infection. These data suggested that one way of LF suppressing SARS-CoV-2 infection is by inhibiting virus to attach to HSPG.

### MUC1, LF and α-LA suppress several existed mutants of SARS-CoV-2 infection and replication

To determine whether MUC1, LF and α-LA still inhibit SARS-CoV-2 variants, we constructed and packaged four SARS-CoV-2 variants of B.1.1.7 (alpha), B.1.351 (beta), P.1 (gamma) and B.1.617.1 (kappa) based on the trVLP system. The Caco-2-N cells were infected with these variants and treated with different concentration of MUC1, rLF and α-LA, respectively (**Figure 8A**). After 48 hours infection, the cells were harvested. Interestingly, MUC1, rLF and α-LA could decrease SARS-CoV-2 variants infection of B.1.1.7 (alpha) (**Figure 8B**), B.1.351 (beta) (**Figure 8C**), P.1 (gamma) (**Figure 8D**) and B.1.617.1 (kappa) (**Figure 8E**) in a dose dependent manner. These results suggested that MUC1, rLF and α-LA still could suppress SARS-CoV-2 variants infection and replication.

**Fig. 8.**
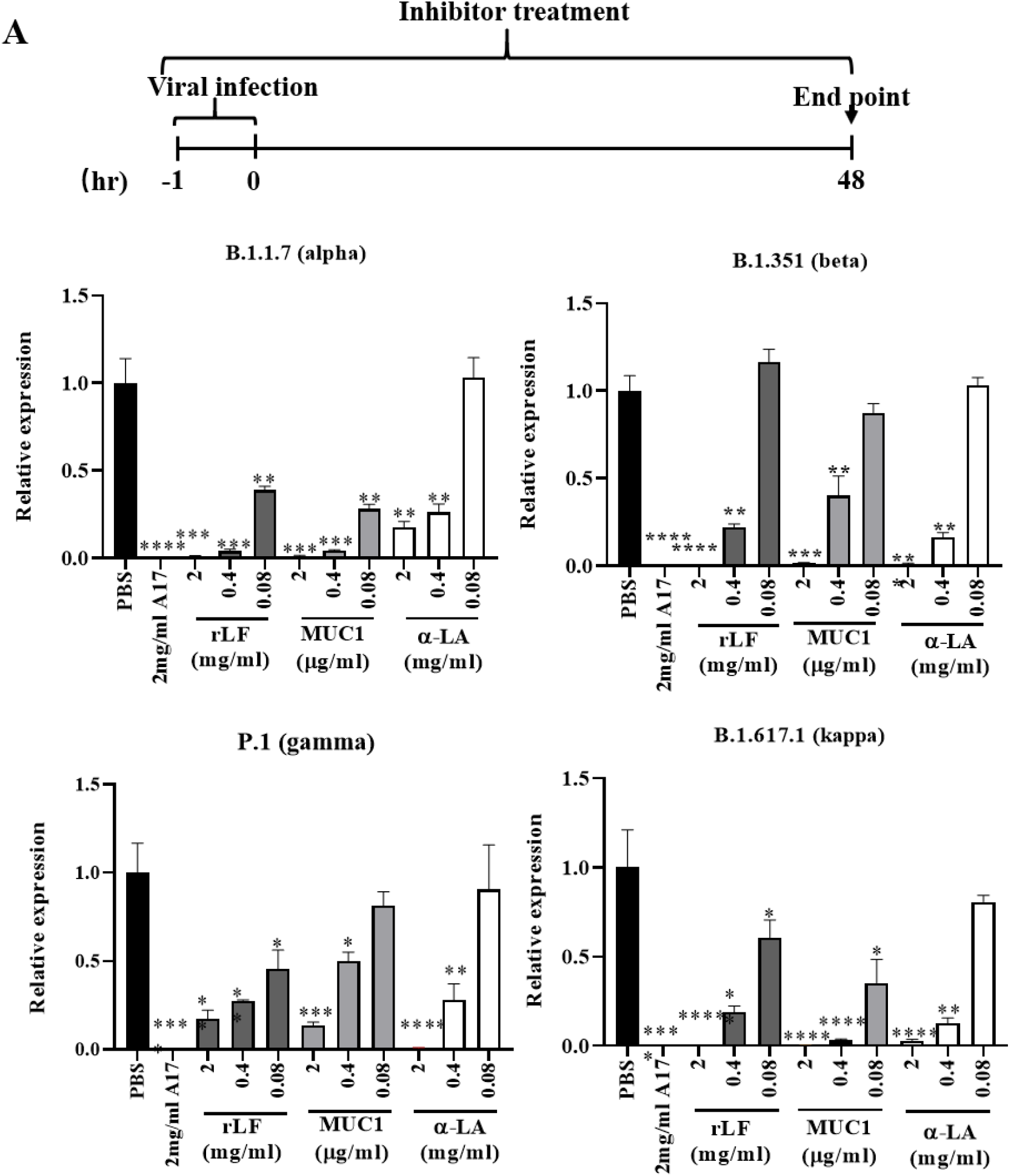
MUC1, LF and α-LA suppress different SARS-CoV-2 variants infection and replication in Caco-2-N cells. (A) Schematic illustration of the treatment experiment of MUC1, LF and α-LA for SARS-CoV-2 variants infection. The caco-2-N cells were seed in the six well plates and infected with SARS-CoV-2 variants of B.1.1.7 (alpha), B.1.351 (beta), P.1 (gamma) and B.1.617.1 (kappa) (MOI=0.1). The cells were treated with MUC1, rLF and α-LA, respectively during the infection. The PBS and the skimmed milk of A17 were used as negative and positive control, respectively. Forty-eight hours post-infection, the cells were harvested and tested. The inhibition analysis of MUC1, rLF and α-LA to SARS-CoV-2 variants of (B) B.1.1.7 (alpha), (C) B.1.351 (beta), (D) P.1 (gamma) and (E) B.1.617.1 (kappa) reflected by intracellular viral RNA were determined by RT-qPCR. Data are presented as mean± SD and repeated at least three times (N = 3), *p<0.05, **p < 0.01, ***p<0.001, ****p<0.0001. α-LA, a-lactalbumin; LF, lactoferrin; A17, skimmed milk.

## Discussion

In this study, we developed an experimental workflow based on LC-MS and SARS-CoV-2 pseudovirus system profiling that allows for effective screening the potential components for inhibiting SARS-CoV-2 infection in vitro. Of these, we identified and confirmed that LF, MUC1 and α-LA in the breastmilk significantly inhibited SARS-CoV-2 infection in the SARS-CoV-2 pseudovirus in Huh7.5 and Vero E6 cells and trVLP infected models in Caco-2-N cells. Our study confirmed the critical role of LF and MUC1 not only in blocking viral attachment and enter to the cells but also in suppressing post-entry replication at multiple cell lines. Moreover, α-LA could only bock viral attachment and entry to the cells. These data shed a light on the groundwork for further antiviral treatment development to SARS-CoV-2 infection and spread.

Although many clinical studies didn’t find solid information of COVID-19 related MTCT through breastfeeding, SARS-CoV-2 positive in breastmilk still confer great concern of the safety of breastfeeding to infants from SARS-CoV-2 infected mothers^14,16,29,30^. Chambers et al. reported that they collected the positive breastmilk from 18 infected women and tested the infectivity in the cell culture system^16^. They didn’t find any viral RNA positive in the experiments, indicating that SARS-CoV-2 in the breastmilk can’t infect and transmit. Consistent to this, our previous published work also showed that even we added the live virus at relative high concentrations in the breastmilk, the virus still could not infect the cells, suggesting that human breastmilk has high quality in anti-SARS-CoV-2 infection^31^. In this study, we also identified LF, MUC1 and α-LA in the milk showed high effective antiviral activity to SARS-CoV-2 infection. Interestingly, the EC_50_ of these factors is high way lower than their concentration in human breastmilk, suggesting the strong antiviral activity of human breastmilk for SARS-CoV-2 infection, which hardly allow the possible live SARS-CoV-2 virus in the milk infect and transmit to the infants during the breastfeeding. For example, the EC_50_ of LF to SARS-CoV-2 is 0.08 mg/ml, while the concentration of LF in human mature breastmilk and colostrum is 1-2 mg/ml and 5-10 mg/ml, respectively. That means, LF shows high concentration in breastmilk and effectively inhibits SARS-CoV-2 possible infection and transmission. This analysis suggested that breastmilk feeding may be safe for infants whose mother are infected with SARS-CoV-2, if the breastmilk was pumped into sterile containers and feed infants with isolation from their mother. However, it still needs further study to confirm the safety of breastfeeding to infants from SARS-CoV-2 infected mother.

SARS-CoV-2 belongs to RNA virus and is easy to mutate during the infection, which resulting in viral escape from antibody neutralization, vaccination, and other drugs targeting the interaction between S protein and ACE-2^4^. Surprisingly, LF, MUC1 and α-LA still could inhibit the existed mutants such as B.1.1.7 (alpha), B.1.351 (beta), P.1 (gamma) and B.1.617.1 (kappa), indicating the strong effects and broad spectrum of these factors for inhibiting SARS-CoV-2 infection.

Most noteworthy, our identified LF, MUC1 and α-LA are safe and available in dietary supplement^32^. All these factors were reported with antiviral activity and their expression was associated with viral infection^24^. Reghunathan R et al. showed that LF expression upregulated during the SARS-CoV infection and blocked viral attachment through HSPG^33^. Moreover, Mirabelli C et al reported that LF screened from the FDA approved drugs also inhibited SASRS-CoV-2 to bind to the cells through competition with HSPG and also modulating innate immune responses to increase expression of interferon stimulated genes and TNFα^21^. Similarly, our study also proved that LF from different species (such as bovine and goat) also inhibited SARS-CoV-2 attachment to HSPG and post-entry replication. LF from human and bovine showed nearly no toxic to all the used cells like Huh7.5, Vero E6 and Caco-2-N cells. However, LF from goat could be toxic to cells at high concentration treatment, indicating that different species has different effects on human cells. Our previous study showed that skimmed milk could suppress the activity of RNA dependent RNA polymerase of SARS-CoV-2^34^. Thus, it still needs further study to analyze the exact mechanism of how LF inhibits SARS-CoV-2 infection and replication.

In addition, MUC1 is an important factor expressing in the lung airway maintaining lung function and health^35^. MUC1 plays important roles in protection of host from infection by pathogens and regulating inflammatory in response to infection^35^. Lu W et al reported that MUC1 expression increased in the airway mucus of critical ill COVID-19 patients^36^. MUC1 sialylated in the surface of cells has the potential to bind influenza A virus (IAV) to reduce viral ability to infection host cells^25^. Consistent to this result, our data showed that MUC1 from milk could inhibit SARS-CoV-2 infection through influence on both viral attachment, entry and post-entry replication. However, affinity assay showed MUC1 didn’t interfere with viral infection based on the ACE-2 target, indicating the possible other mechanism involved in the infection. Interestingly, we used the recombined MUC1 C-terminal unit (MUC1-C) without any variable number tandem repeat (VNTR) region also showed high inhibitory activity to SARS-CoV-2 infection, suggesting the possible other mechanisms for MUC1 to inhibit SARS-CoV-2 infection and replication. It has been demonstrated by many studies that MUC1 play an important role in inhibiting Toll like receptor and NOD-like receptor protein 3 dependent inflammation^35^. These results suggested that MUC1 could be used as a potential drug for inhibiting SARS-CoV-2 infection and related inflammation to reduce the severity of COVID-19. However, it needs more further studies to support it. As for α-LA, it seems that α-LA could only block viral attachment to the cells through interfere viral binding to ACE-2 receptor.

Collectively, we identified several factors from human breastmilk (LF, MUC1 and α-LA) effectively inhibit SARS-CoV-2 and their variants infection and replication by viral attachment, entry and post-entry replication. Although our findings are promising and showed its potential role in the safety of breastmilk feeding in the clinic, further studies are still needed to confirm our findings in animal and clinical studies and further modification of these factors molecular structure for clinical usage.

## Resource Availability

### Lead Contact

Further information and requests for resources and reagents should be directed to and will be fulfilled by the Lead Contact, Kuanhui Xiang

### Materials Availability

All materials in this study are available from the Lead Contact with a completed Materials Transfer Agreement.

### Data and Code Availability

The original sequencing datasets for trVLP of SARS-CoV-2 can be found on the Wuhan-Hu-1, MN908947

### Experimental model and subject details

#### Cell lines and key reagents

HEK293T, Vero E6, Huh7.5 and Caco-2-N cells were maintained in Dulbecco’s modified Eagle medium (DMEM, Gibco, China) supplemented with 10% fetal bovine serum (FBS) in a humidified 5% (vol/col) CO_2_ incubator at 37°C. All cell lines were tested negative for mycoplasma. Goat and cow whey proteins were purchased from Sigma (USA). The recombined lactoferrin (rLF), human lactoferrin (hLF) and bovine lactoferrin (bLF) were purchased from Sigma (USA). Lactadherin, a-lactalbumin were purchased from Sigma (USA). MUC1 encoding from Ala23 to Gly167 is expressed with a Fc tag at the C-terminus and purchased from Novoprotein (China).

#### Viruses

SARS-CoV-2 pseudovirus with luciferase and GFP expression was kindly shared by Prof. Youchun Wang (National Institutes for Food and Drug Control, China) and Prof. Ningshao Xia (Xiamen University), respectively. The trVLP and their mutants of B.1.1.7 (alpha), B.1.351 (beta), P.1 (gamma) and B.1.617.1 (kappa) expressing GFP replacing viral nucleocapsid gene (N) were made as previously described^7^. Caco-2-N cells were infected with the trVLP to amplify viruses^28^. P5 to P10 were used in this study.

#### Skimmed milk samples preparation

The skimmed milk samples preparation was performed as previously described. Briefly, the milk samples were centrifuged for 15 min at 4,000 x g at 4°C and the lower aqueous phase was collected and used for further experiments and analysis^34^.

#### The trVLP virus production

The viral production was performed as previously described^28^. Briefly, the viral RNA transcript and viral N mRNA were obtained by in vitro transcribed. Viral RNA and N mRNA were mixed and electroporated into Caco-2-N cells. The produced virus was collected and amplified in Caco-2-N cells for several passages (to P10).

#### SARS-CoV-2 pseudovirus and trVLP infection assay

SARS-CoV-2 pseudovirus infection was performed as described^34^. The cells were infected with viral inocula of 650 TCID_50_/well. One day post infection (1dpi), the cells were lysed and the luminescence was measured according to the manufacture’s protocol. For trVLP, Caco-2-N cells were infected trVLP at multiplicity of infection (MOI) of 0.1 as described. Several days post-infection, the mRNA levels of trVLP and GAPDH were determined by RT-qPCR.

#### In-gel digestion and LC-MS/MS analysis

Upon SDS-PAGE fractionation, the band of interest was excised and subjected to in-gel trypsin digestion as previously described^27^. LC-MS analyses of protein digests were carried out on a hybrid ion trap-Orbitrap mass spectrometer (LTQ Orbitrap Velos, Thermo Scientific) coupled with nanoflow reversed-phase liquid chromatography (EASY-nLC 1000, Thermo Scientific). The capillary column (75 μm× 150 mm) with a laser-pulled electrospray tip (Model P-2000, Sutter instruments) was home-packed with 4 μm, 100 Å Magic C18AQ silicabased particles (Michrom BioResources Inc., Auburn, CA) and run at 250 nL/min with the following mobile phases (A: 97% water, 3% acetonitrile, and 0.1% formic acid; B: 80% acetonitrile, 20% water, and 0.1% formic acid). The LC gradient started at 7% B for 3 min and then was linearly increased to 37% in 40 min. Next, the gradient was quickly ramped to 90% in 2 min and stayed there for 10 min. Eluted peptides from the capillary column were electrosprayed directly onto the mass spectrometer for MS and MS/MS analyses in a data dependent acquisition mode. One full MS scan (m/z 400–1200) was acquired by the Orbitrap mass analyzer with R = 60,000 and simultaneously the ten most intense ions were selected for fragmentation under collisioninduced dissociation (CID). Dynamic exclusion was set with repeat duration of 30 s and exclusion duration of 12 s.

#### Skimmed milk separation

Skimmed milk was precipitated by ammonium sulfate hydrochloride, after centrifugation at 15000×g for 2 h, samples were applied to a HiTrap SP HP cation exchange chromatography column (GE Healthcare) equilibrated with buffer A (20 mM Phosphate Buffer, pH7.2). The column was washed with three column vol of buffer A, and bound material was eluted with 10 column vol of buffer B (20 mM Phosphate Buffer 1M NaCl, pH7.2), flow-through and eluted fractions was assayed for testing and identifying the exact fraction with inhibition capacity of SARS-CoV-2 pseudovirus infection. Then, the fraction with capacity to inhibit viral infection was applied to a HiTrap Q HP anion exchange chromatography column (GE Healthcare) equilibrated with buffer A (20 mM Phosphate Buffer, pH7.2). The column was washed with three column vol of buffer A, and bound material was eluted with 10 column vol of buffer B (20 mM Phosphate Buffer 1M NaCl, pH7.2), flow-through and eluted fractions was assayed for testing and identifying the exact fraction with inhibition capacity of SARS-CoV-2 pseudovirus infection. The effective fraction was mixed with 1% NP-40 and performed to a HiTrap SP HP anion exchange chromatography column (GE Healthcare) equilibrated with buffer A (20 mM Phosphate Buffer, pH7.2). The column was washed with three column vol of buffer A, and bound material was eluted with 10 column vol of buffer B (20 mM Phosphate Buffer 1M NaCl, pH7.2), flow-through and eluted fractions was assayed for testing and identifying the exact fraction with inhibition capacity of SARS-CoV-2 pseudovirus infection. Afterwards, the effective fraction of peak 1 was applied to Superdex 75 10/300 GL size-exclusion chromatography column quilibrated in buffer A, fraction was collected for In-gel digestion and LC-MS/MS analysis.

#### Viral attachment assay

Caco-2N cells were seed at the 24-well plates with 80,000 cells/well one day before the infection. LF (2 mg/ml), MUC1 (2 ug/ml) and α-LA (2 mg/ml) was mixed with trVLP (MOI=1) at 4°C for 1h, respectively. The mixture was added into the cells and put at 4°C for 2h to allow viral attachment to cells. After washing out of free virus, cell surface GX_P2V was extracted and quantified by RT-qPCR. For SARS-CoV-2 pseudovirus, Huh7.5 and Vero E6 cells were seed at the 96-well plates with 20,000 cells/well one day before infection. LF (2 mg/ml), MUC1 (2 ug/ml) and α-LA (2 mg/ml) was mixed with trVLP (MOI=1) at 4°C for 1h, respectively. The mixture was added into the cells and put at 4°C for 2h to allow viral attachment to cells. After washing out of free virus, the cells were incubated at 37°C for 24 h. The luciferase assay was performed to detect SARS-CoV-2 pseudovirus infection. For SARS-CoV-2 pseudovirus, the method was described as previously described^34^.

#### Viral entry assay

Caco-2-N cells were exposed to trVLP (MOI=1) at 4 °C for 1h. Then, the cells were washed with PBS for 3 times. LF (2 mg/ml), MUC1 (2 ug/ml) and α-LA (2 mg/ml) were added into the media and incubated at 37°C for 1h to allow viral internalization into cells, respectively. The mRNA of trVLP was measured by RT-qPCR. GFP were detected by a fluorescence microscope (ECHO laboratories, USA). For SARS-CoV-2 pseudovirus, the method was described as previously described^34^.

#### Viral post-entry assay

Caco-2-N cells were infected with trVLP and incubated at 37 °C for 1h. After washing out of the free viruses, the cells were cultured in the media containing LF (2 mg/ml), MUC1 (2 ug/ml) and α-LA (2 mg/ml), respectively. Different time point of post-infection, the mRNA of trVLP was measured by RT-qPCR. GFP were detected by a fluorescence microscope (ECHO laboratories, USA). For SARS-CoV-2 pseudovirus, the method was described as previously described^34^.

#### Viral RNA extraction and quantification

The RNA extraction and quantification were performed as previously described^28^. Briefly, Total RNA of cells was isolated with the RNAprep pure Cell Culture/Bacterial total RNA extraction kit (Tiangen biotech. Co., China). The cDNA was synthesized by RevertAid first strand cDNA synthesis kit (Invitrogen, USA). The qPCR for SARS-CoV-2 RNA was performed as previously described. The qPCR primers for viral RNA were as follows: THU-2190 (5’-CGAAAGGTAAGATGGAGAGCC-3’) and THU-2191 (5’-TGTTGACGTGCCTCTGATAAG-3’). GAPDH was used to normalize all the data^28^.

#### Western blotting

Western blotting was performed as described previously^34^. Briefly, the cells samples were loaded on the 12% SDS-PAGE gel and transferred to a polyvinylidene fluoride membrane. The antibody of anti-SARS-CoV-2 S protein against spike protein and anti-β-actin were used at 1:2000 dilutions. The second antibody of HRP-conjugated affinipure Goat anti-mouse IgG (H+L) were diluted at 1:20000. SuperSignal® West Femto Maximum Sensitivity Chemiluminescent Substrate (Thermo Scientific, USA) was used for imaging.

#### Affinity assay between ACE2 and SARS-CoV-2 Spike protein

The influence of LF, MUC1 and α-LA on the affinity between ACE2 and SARS-CoV-2 S RBD was performed as previously described^34^. Briefly, the ACE-2 (Novoprotein, China) was immobilized on the MaxiSORP ELISA plate at 100ng per well in 50 μl of 100 mM carbonate-bicarbonate coating buffer over night at 4 °C. HRP-conjugated SARS-CoV-2 RBD (Novoprotein, China) at final concentration of 1 ng/μl was mixed with serially diluted human breastmilk. LF, MUC1 and α-LA at different concentrations were added into the ACE-2-coated plate for 1h at room temperature, respectively. The absorbance reading at 450 nm and 570 nm were acquired using the Cytation 5 microplate reader (Bio Tek).

#### Statistical analysis

Statistical analyses were analyzed using GraphPad Prism 8 software (GraphPad Software Inc., San Diego, CA, USA). Data are presented as mean± SD and repeated at least three times (N = 3). Comparisons between the two groups were analyzed using the Student’s t tests. Values of p<0.05 was considered statistically significant.

## Acknowledgements

We gratefully thank Prof. Youchun Wang and Weijin Huang (National Institutes for Food and Drug Control, China) for sharing the luciferase reported pseudovirus of SARS-CoV-2 and Ningshao Xia (Xiamen University, China) for sharing the GFP reported pseudovirus of SARS-CoV-2. We thank Prof. Xiaodong Su (Peking University) for helpful discussion.

