## supplemental figures for "Inhibition of SAR S-CoV-2 infection and replication by lactoferrin, MUC1 and α-lactalbumin identified in human breastmilk"

**Supplementary figures and figure legends**

```
  1 MKLVFLVLLF LGALGLCLAG RRRSVQWCAV SQPEATKCFQ WQRNMRKVRG
 51 PPVSCIKRDS PIQCIQAIAE NRADAVTLDG GFIYEAGLAP YKLRPVAAEV
101 YGTERQPRTH YYAVAVVKKG GSFQLNELQG LKSCHTGLRR TAGWNVPIGT
151 LRPFLNWTGP PEPIEAAVAR FFSASCVPGA DKGQFPNLCR LCAGTGENKC
201 AFSSQEPYFS YSGAFKCLRD GAGDVAFIRE STVFEDLSDE AERDEYELLC
251 PDNTRKPVDK FKDCHLARVP SHAVVARSVN GKEDAIWNLL RQAQEKFGKD
301 KSPKFQLFGS PSGQKDLLFK DSAIGFSRVP PRIDSGLYLG SGYFTAIQNL
351 RKSEEEVAAR RARVVWCAVG EQELRKCNQW SGLSEGSVTC SSASTTEDCI
401 ALVLKGEADA MSLDGGYVYT AGKCGLVPVL AENYKSQQSS DPDPNCVDRP
451 VEGYLAVAVV RRSDTSLTWN SVKGKKSCHT AVDRTAGWNI PMGLLFNQTG
501 SCKFDEYFSQ SCAPGSDPRS NLCALCIGDE QGENKCVPNS NERYYGYTGA
551 FRCLAENAGD VAFVKDVTVL QNTDGNNNEA WAKDLKLADF ALLCLDGKRK
601 PVTEARSCHL AMAPNHAVVS RMDKVERLKQ VLLHQQAKFG RNGSDCPDKF
651 CLFQSETKNL LFNDNTECLA RLHGKTTYEK YLGPQYVAGI TNLKKCSTSP
701 LLEACEFLRK
```

**Figure S1. Lactotransferrin eluate from Anion-Exchange Chromatography, sequence coverage (32%) of mascot search (matched peptides shown in red)**

```
  1 MKLVFLVLLF LGALGLCLAG RRRSVQWCAV SQPEATKCFQ WQRNMRKVRG
 51 PPVSCIKRDS PIQCIQAIAE NRADAVTLDG GFIYEAGLAP YKLRPVAAEV
101 YGTERQPRTH YYAVAVVKKG GSFQLNELQG LKSCHTGLRR TAGWNVPIGT
151 LRPFLNWTGP PEPIEAAVAR FFSASCVPGA DKGQFPNLCR LCAGTGENKC
201 AFSSQEPYFS YSGAFKCLRD GAGDVAFIRE STVFEDLSDE AERDEYELLC
251 PDNTRKPVDK FKDCHLARVP SHAVVARSVN GKEDAIWNLL RQAQEKFGKD
301 KSPKFQLFGS PSGQKDLLFK DSAIGFSRVP PRIDSGLYLG SGYFTAIQNL
351 RKSEEEVAAR RARVVWCAVG EQELRKCNQW SGLSEGSVTC SSASTTEDCI
401 ALVLKGEADA MSLDGGYVYT AGKCGLVPVL AENYKSQQSS DPDPNCVDRP
451 VEGYLAVAVV RRSDTSLTWN SVKGKKSCHT AVDRTAGWNI PMGLLFNQTG
501 SCKFDEYFSQ SCAPGSDPRS NLCALCIGDE QGENKCVPNS NERYYGYTGA
551 FRCLAENAGD VAFVKDVTVL QNTDGNNNEA WAKDLKLADF ALLCLDGKRK
601 PVTEARSCHL AMAPNHAVVS RMDKVERLKQ VLLHQQAKFG RNGSDCPDKF
651 CLFQSETKNL LFNDNTECLA RLHGKTTYEK YLGPQYVAGI TNLKKCSTSP
701 LLEACEFLRK
```

**Figure S2. Lactotransferrin eluate from Size-Exclusion Chromatography, sequence coverage (30%) of mascot search (matched peptides shown in red)**

```
  1 MKLVFLVLLF LGALGLCLAG RRRSVQWCAV SQPEATKCFQ WQRNMRKVRG
 51 PPVSCIKRDS PIQCIQAIAE NRADAVTLDG GFIYEAGLAP YKLRPVAAEV
101 YGTERQPRTH YYAVAVVKKG GSFQLNELQG LKSCHTGLRR TAGWNVPIGT
151 LRPFLNWTGP PEPIEAAVAR FFSASCVPGA DKGQFPNLCR LCAGTGENKC
201 AFSSQEPYFS YSGAFKCLRD GAGDVAFIRE STVFEDLSDE AERDEYELLC
251 PDNTRKPVDK FKDCHLARVP SHAVVARSVN GKEDAIWNLL RQAQEKFGKD
301 KSPKFQLFGS PSGQKDLLFK DSAIGFSRVP PRIDSGLYLG SGYFTAIQNL
351 RKSEEEVAAR RARVVWCAVG EQELRKCNQW SGLSEGSVTC SSASTTEDCI
401 ALVLKGEADA MSLDGGYVYT AGKCGLVPVL AENYKSQQSS DPDPNCVDRP
451 VEGYLAVAVV RRSDTSLTWN SVKGKKSCHT AVDRTAGWNI PMGLLFNQTG
501 SCKFDEYFSQ SCAPGSDPRS NLCALCIGDE QGENKCVPNS NERYYGYTGA
551 FRCLAENAGD VAFVKDVTVL QNTDGNNNEA WAKDLKLADF ALLCLDGKRK
601 PVTEARSCHL AMAPNHAVVS RMDKVERLKQ VLLHQQAKFG RNGSDCPDKF
651 CLFQSETKNL LFNDNTECLA RLHGKTTYEK YLGPQYVAGI TNLKKCSTSP
701 LLEACEFLRK
```

**Figure S3. Lactotransferrin eluate from Cation-Exchange Chromatography, sequence coverage (26%) of mascot search (matched peptides shown in red)**

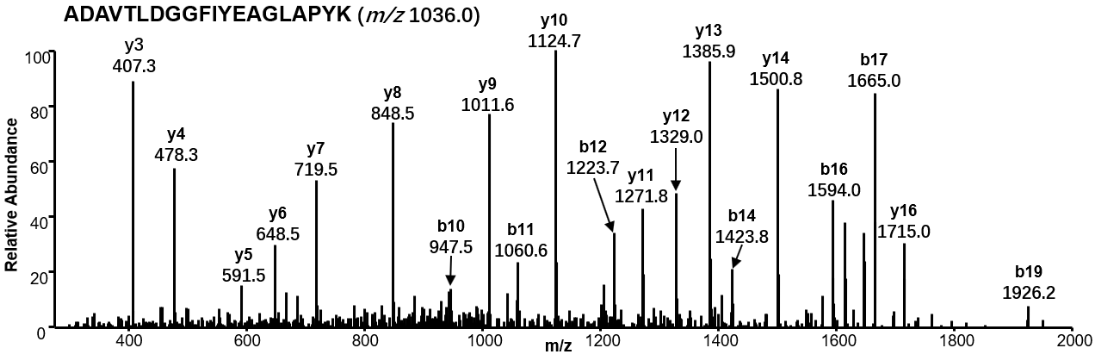


**Figure S4. Unique peptide of lactotransferrin identification by tandem mass spectrometry (MS/MS) analysis.**

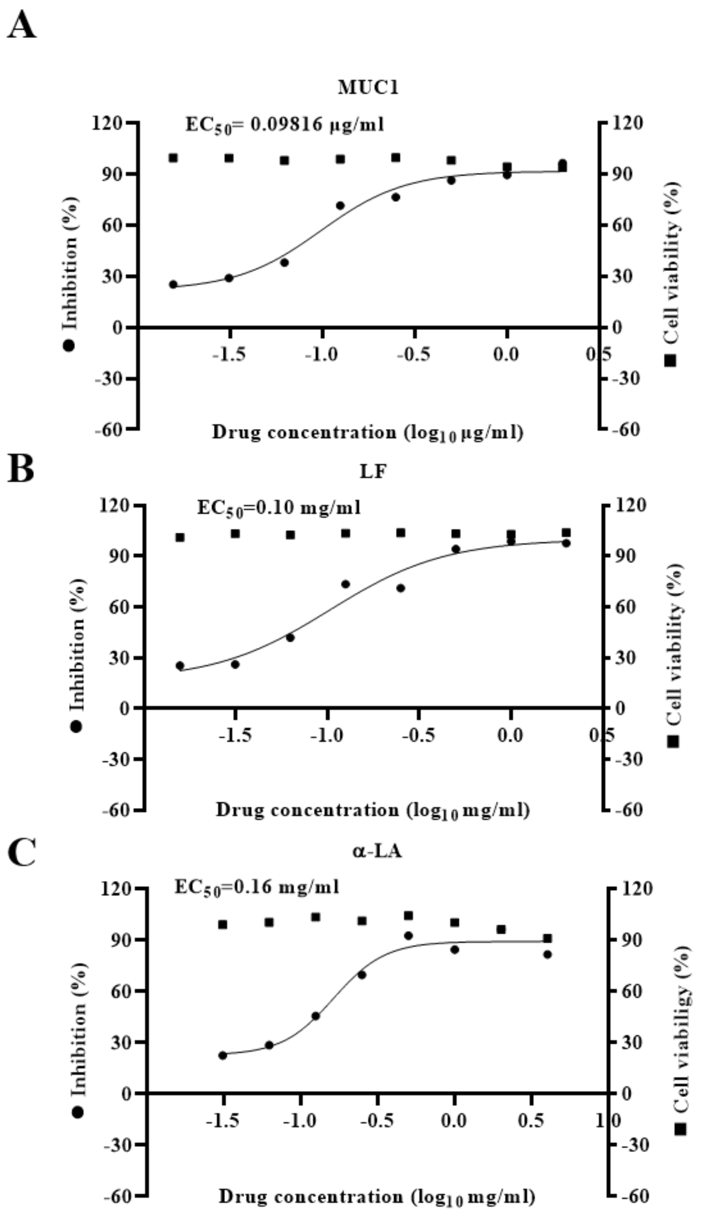


**Figure S5. MUC1, LF and α-LA treatment dose-dependently suppressed viral infection with reduced luminescence in SARS-CoV-2 pseudovirus infected Vero E6 cells.** The inhibition analysis reflected by luminescence of SARS-CoV-2 pseudovirus (650 TCID50/cell) infected Vero E6 cells treated with serial doses of MUC1 (A), LF (B)and α-LA (C) were determined by luminescence according to the manufacture's instruction.

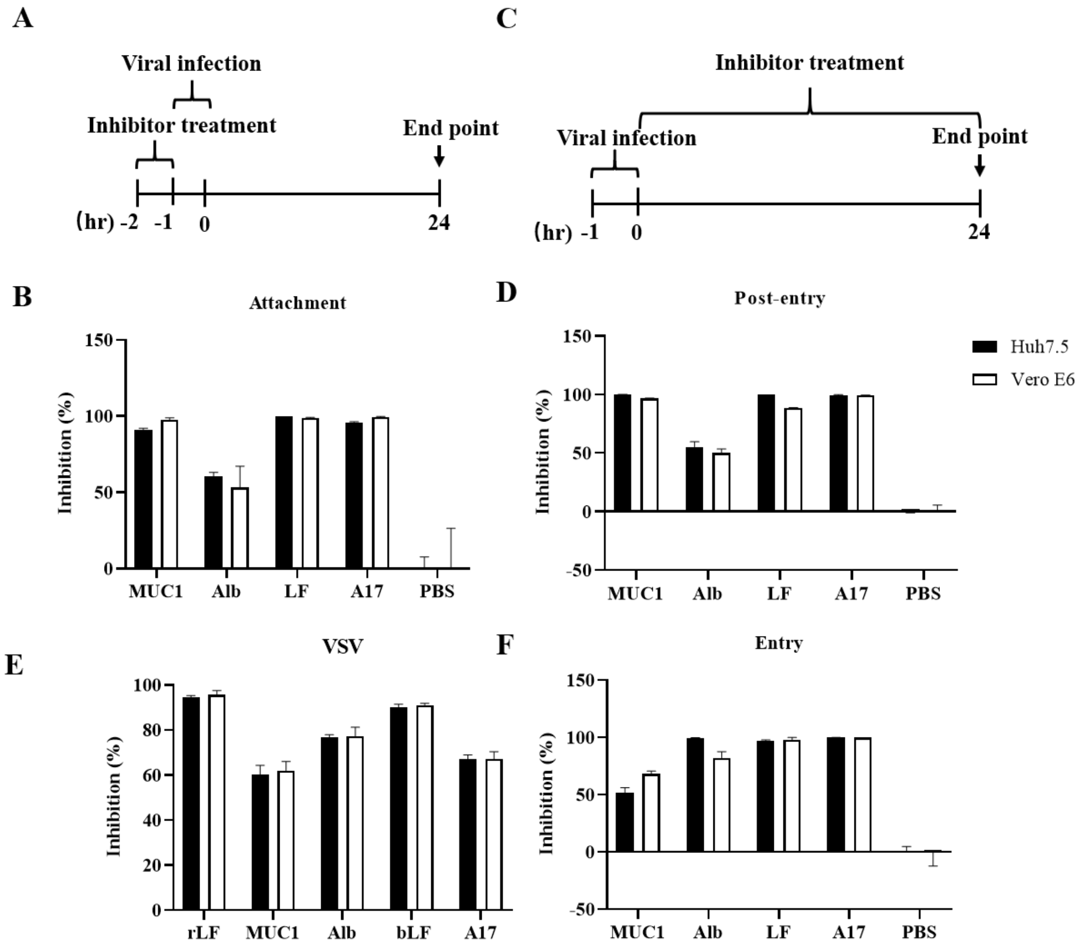


**Figure S6. MUC1, LF and α-LA treatment affects all stages of viral infection including attachment, entry and post-entry in SARS-CoV-2 infected Huh7.5 and Vero E6 cells.** (A) schematic illustration of the pretreatment experiment. (B) Inhibition analysis of MUC1, α-LA and LF to SARS-CoV-2 pseudovirus infection. The cells were lysed and the luminescence was detected according to the manufacture's instruction 24 hours post-infection. (C) Schematic illustration of the post-infection treatment experiment. (D) The luminescence was detected to determine inhibition of SARS-CoV-2 pseudovirus infection. (E) The luminescence was detected to determine inhibition of VSV post-infection. (F) Inhibition analysis of MUC1, α-LA and LF to viral entry stage. The SARS-CoV-2 pseudovirus was mixed with Huh7.5 or Vero E6 cells at 4 °C for 1 hour then discarded the supernatant and washed with PBS for three times. The cells were then added with fresh media with MUC1, α-LA, rLF, hLF, bLF and A17 treatment at 37 °C for 2 hours. After that, the supernatant was disregarded and the cells were reloaded with fresh media and cultured for 24 hours. The cells were lysed and and the luminescence was detected according to the manufacture's instruction 24 hours post-infection. MUC1 (2 ug/ml), α-LA (a-lactalbumin, 2 mg/ml), rLF (recombinant lactoferrin, 2 mg/ml), hLF (human lactoferrin, 2 mg/ml), bLF (bovine lactoferrin, 2 mg/ml) and A17 (skimmed milk, 2 mg/ml). Data are presented as mean± SD and repeated at least three times (N = 3).

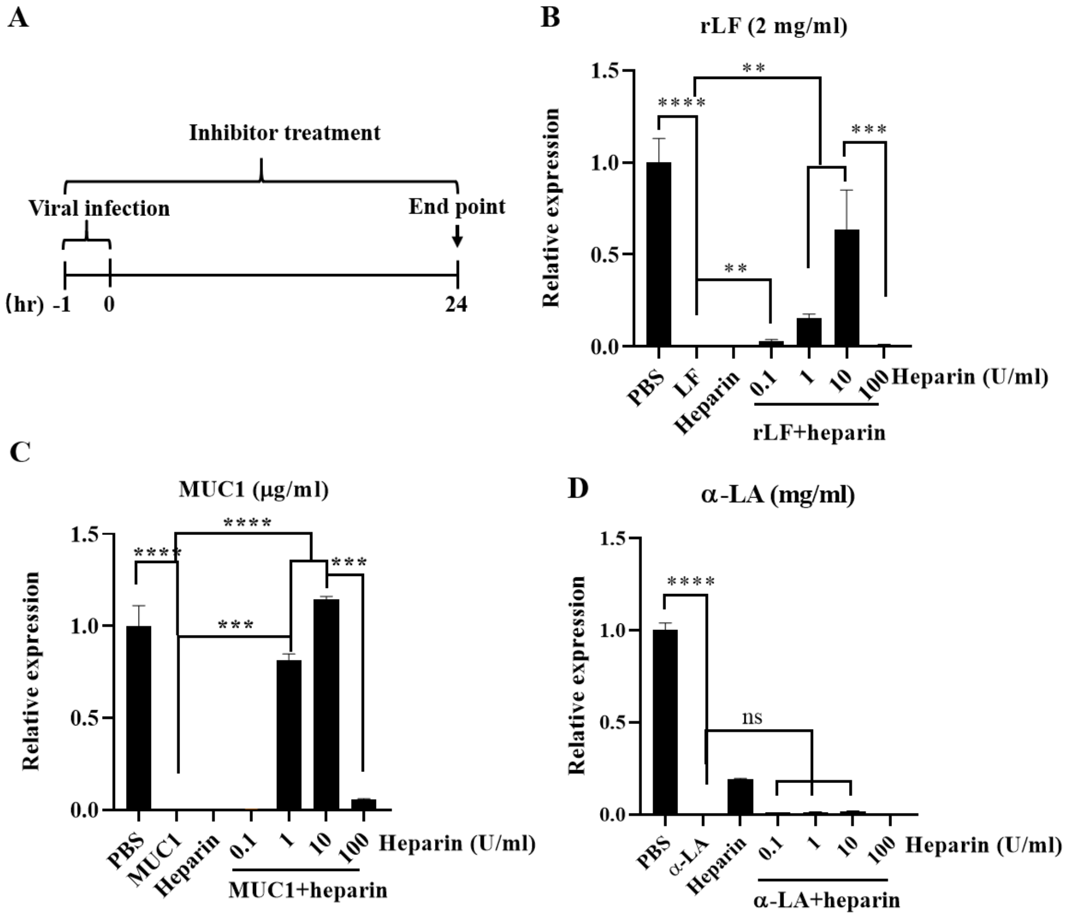


**Fig. S7 MUC1 and LF inhibit SARS-CoV-2 attaching to HSPG in the trVLP infected Caco-2-N cell system**. (A) Schematic illustration of experiment design. Inhibition of SARS-CoV-2 trVLP infection in Caco-2-N cells were treated with heparin for 10 min at the dosage of 0.1 U/ml, 1 U/ml, 10 U/ml and 100 U/ml. Then, 2 mg/ml LF, 2 ug/ml of MUC1 and 2 mg/ml of a-LA were added to each group and incubated at 37°C for 1h. RT-qPCR analysis was performed to detect SARS-CoV-2 RNA levels. Data are presented as mean± SD and repeated at least three times (N = 3), $^{**}p < 0.01$, $^{***}p < 0.001$, $^{****}p<0.0001$. HSPG: heparan sulfate proteoglycans; LF: lactoferrin; MUC1: mucin 1; a-LA: a-lactalbumin; ns: no significant.
